# The zebrafish presomitic mesoderm elongates through compression-extension

**DOI:** 10.1101/2021.03.11.434927

**Authors:** Lewis Thomson, Leila Muresan, Benjamin Steventon

## Abstract

In vertebrate embryos the presomitic mesoderm become progressively segmented into somites at the anterior end while extending along the anterior-posterior axis. A commonly adopted model to explain how this tissue elongates is that of posterior growth, driven in part by the addition of new cells from uncommitted progenitor populations in the tailbud. However, in zebrafish, much of somitogenesis is associated with an absence of overall volume increase and posterior progenitors do not contribute new cells until the final stages of somitogenesis. Here, we perform a comprehensive 3D morphometric analysis of the paraxial mesoderm and reveal that extension is linked to a volumetric decrease, compression in both dorsal-ventral and medio-lateral axes, and an increase in cell density. We also find that individual cells decrease in their cell volume over successive somite stages. Live cell tracking confirms that much of this tissue deformation occurs within the presomitic mesoderm progenitor zone and is associated with non-directional rearrangement. Furthermore, unlike the trunk somites that are laid down during gastrulation, tail somites develop from a tissue that can continue to elongate in the absence of functional PCP signalling. Taken together, we propose a compression-extension mechanism of tissue elongation that highlights the need to better understand the role of tissue intrinsic and extrinsic forces play in regulating morphogenesis.

## 1. Introduction

A key process in early development is the progressive elongation of the embryo along the anterior-posterior axis. In vertebrates, this is coupled with the segmentation of the paraxial mesoderm into somites, that occurs in a clock-like manner from the anterior to the posterior as development continues (Cooke and Zeeman, 1976; Dequéant and Pourquié, 2008; Richardson et al., 1998). To ensure that the appropriate number of somites are generated upon completion of somitogenesis, this process must be balanced with the elongation of the presomitic mesoderm (PSM) (Gomez et al., 2008; Gomez and Pourquie, 2009). A dominant model for PSM elongation is that of posterior growth, based on the idea that cells are continually being added from progenitor populations in the tailbud and the widespread conservation of a common set of gene regulatory interactions in this process (Martin and Kimelman, 2009). However, fast-developing organisms such as zebrafish elongate their body axis in the absence of volumetric growth at the posterior end of the embryo (Bouldin et al., 2014; Steventon et al., 2016; Zhang et al., 2008). In addition, the tailbud population of neuromesodermal progenitors delay their contribution to the PSM until late stages of somitogenesis (Attardi et al., 2018). Therefore, it remains unclear as to how the zebrafish PSM elongations in the absence of posterior growth and progenitor addition.

Several recent studies have highlighted the importance of regulating the cell movements and fluidity of the PSM as cells leave the posterior progenitor domain in the tailbud (expressing *msgn1* (Fior et al., 2012)) and enter the PSM proper (expressing *tbx6* (Griffin et al., 1998)). We refer to these regions as the posterior PSM and the anterior PSM, respectively, given that they are two regions of one continuous tissue, rather than separate tissues. Directional variability in the posterior PSM has been shown to be important for facilitating elongation by ensuring that the posterior flow of dorsal cells into this region is translated into a continuous, bilaterally symmetrical contribution to the anterior PSM (Lawton et al., 2013). Similarly, Mongera et al. (2018) showed that there is a gradient of yield stress from posterior to anterior; with the posterior PSM being more “fluid-like” and showing more cell mixing than the anterior PSM. These authors proposed, using a 2D modelling approach, that a jamming transition between these two regions ensures that cell addition into the posterior PSM is translated into unidirectional elongation, rather than isotropic expansion. Importantly, both explanations were based on experiments using mid-somitogenesis embryos (10-14SS), in which cells are still being added to the posterior PSM from the dorsal medial zone through gastrulation movements. How the PSM continues to elongate at later somite stages has not yet been fully determined.

An alternative explanation is that posterior PSM cells are not moving randomly but migrating anteriorly; competing with each other to move out of the tailbud and into the anterior PSM (Manning and Kimelman, 2015). However, given that the tissue is elongating, defining cell movement as anterior vs posterior is difficult, and will depend on the frame of reference. Presumably, in an elongating tissue, cells relative to the posterior end will all move anteriorly - but relative to the anterior end will move posteriorly. To answer the question of how the paraxial mesoderm elongates in the absence of posterior cell addition and growth, a long-term, multi-scale, threedimensional approach is required. Through long-term 3D morphometrics and celltracking we show that elongation of the PSM is associated with overall tissue compression. However, this “compression-extension” is not driven by local cell intercalation, but through convergence of cells to the midline coupled with non-orientated cell intercalation.

## 2. Materials and methods

### 2.1. Animal husbandry and microinjection

Zebrafish embryos were raised in standard E3 media at 28°C. Wild type lines used were Tüpfel Long Fin (TL), AB, and AB/TL. The following transgenic lines were also used: *tbx16::GFP* (Wells et al., 2011) *, h2a::mCherry*, and *h2b::GFP*. Embryos staged according to Kimmel et al. (1995). mRNA (KikGR and DEP+) was recovered from *E. coli* plasmid stocks using standard protocols and diluted to 100-250 ng/μl in nuclease-free water, with phenol red added (5%) to help with visibility during microinjection. Embryos were microinjected (using pulled capillary needles) at the 1-cell stage.

### 2.2. in situ hybridization chain reaction (HCR)

Embryos were manually dechorionated and fixed in 4% PFA in PBS (without calcium and magnesium) overnight at 4°C, then dehydrated in methanol and stored at −20°C. HCR was then performed using standard zebrafish protocol (Choi et al., 2018) and nuclei were stained with DAPI. The dehydration step was omitted when also staining membranes with phalloidin.

### 2.3. Imaging

For imaging fixed embryos, the tail was cut off and mounted in 80% glycerol. Live embryos were anaesthetised with tricaine and mounted whole in LMP agarose (1% in E3 media), covered in E3 media. The posterior part of the embryo was freed from agarose by cutting away the agarose with a microinjection needle, as described by Hirsinger and Steventon, (2017), which allowed normal development of the trunk and tail. All confocal imaging was performed on a Zeiss LSM 700 (inverted) and a Leica SP8 (inverted). Imaging was performed using 10x (air), 20x (air), or 40x (oil) objectives depending on the resolution required for analysis and on embryo size/mounting method. Live confocal imaging was performed with the use of a heated chamber (heated to 28°C). Live two-photon imaging was performed on a two-photon microscope (upright) with a 25x (water) objective and a heated chamber (heated to 28°C).

### 2.4. Image analysis

#### Imaris surfaces creation

3D tissue reconstructions of paraxial mesoderm tissues (the PSM and nascent somite for each stage) were generated by creating “surfaces” in Imaris. This involved manually drawing “contours” around the tissue at regular z-intervals, which were then automatically translated into a 3D surface (Figure 1B; Figure S1A-C; Movie S1). Surfaces were created for one lateral half of the PSM, and one of the nascent somite pair, at each stage. Imaris surfaces provided volumes of paraxial mesoderm tissues over time. A separate automatic surface was also generated of the posterior PSM/progenitor region, using *msgn1* HCR signal. This was useful for taking separate dimension measurements of the posterior and anterior PSM (see below).

**Figure 1:**
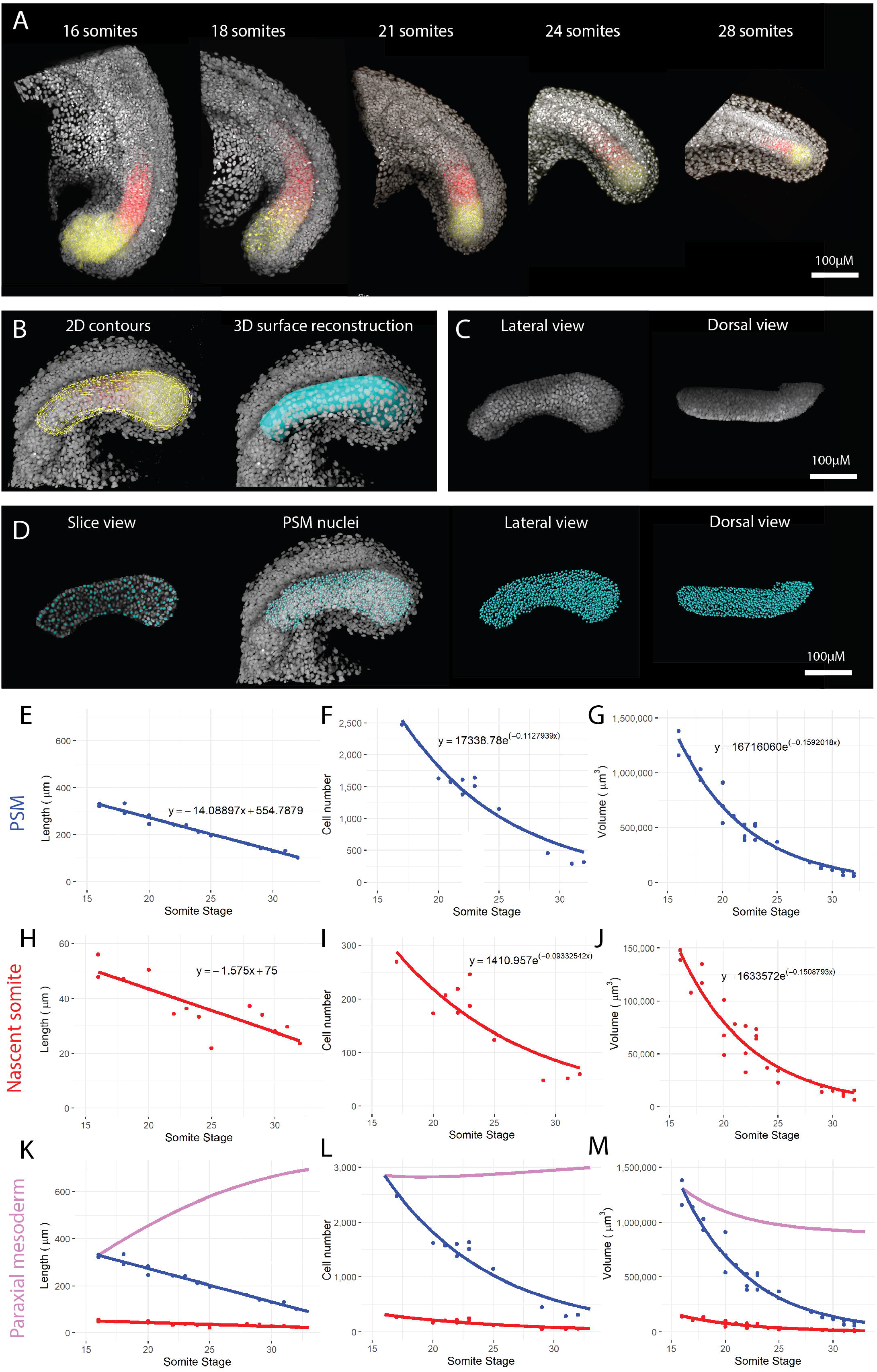
PSM elongation occurs in the absence of growth. (A) *in situ* hybridization chain reaction (HCR) was used to stain for *msgn1* (yellow) and *tbx6* (red), markers of the posterior and anterior PSM, respectively, from the 16 somite-stage to the end of somitogenesis. Nuclei were stained with DAPI (grey). Images are maximum projections of confocal stacks. (B) 2D contours (yellow outlines, left image) were manually drawn around the PSM at regular z-slices, up to the midline, to generate 3D surface reconstructions of the PSM (cyan, right image) and the nascent somite (not shown) at each stage. (C) DAPI signal was isolated from each surface to show only nuclei in that tissue. (D) Spots were generated, marking the centre of each isolated nucleus (shown in slice view (left image) and 3D view (other images)). The length (E), cell number (F), and volume (G) of the PSM at each stage was measured. Possible trendline equations for each set of measurements were calculated in R, and AIC was used to determine the best statistical model (linear, exponential, quadratic, etc.).The same was done for the length (H), cell number (I), and volume (J) of each somite at its time of formation. These trendline equations were then used to calculate paraxial mesoderm values for length (K), cell number (L), and volume (M).

#### Imaris spots creation

DAPI signal was isolated from the surfaces and used to generate a “spot” of each nucleus in paraxial mesoderm tissues. The estimated cell diameter used was 4 μm, and the minimum fluorescence intensity threshold was set to 0. These parameters gave the most accurate cell number estimates. This accuracy was validated by “masking” spots from the nuclei (to create a small black hole in each nucleus), then creating a new channel (given a different colour) with only DAPI signal inside the spots (to create a coloured dot in each nucleus). This method allowed us to check individual z-slices for the presence of one coloured dot in each nucleus. Estimated cell diameters below 4 μm led to multiple dots per nucleus (i.e. overestimation of cell number), while estimate cell diameters above 4 μm led to many nuclei with no dots (i.e. underestimation of cell number), as did setting any minimum fluorescence intensity threshold. Imaris spots provided cell numbers and 3D positions for paraxial mesoderm tissues over time.

#### Dimension measurements

This was done manually in Imaris using “measurement points” and the 3D surfaces (described above). To measure length (anteroposterior (AP) axis) of the PSM, both the manual PSM surface and the automatic *msgn1* surface were used. The measurement was taken through the middle of the tissue, following the tissue curve by taking two measurements: one from the posterior medial face to the anterior face of the *msgn1* surface, and one from here to the anterior face of the PSM surface. This length measurement thus considered the tissue curve, as well as the fact that the surface is only one lateral half of the true tissue. For both width (mediolateral (ML) axis) and height (dorsoventral (DV) axis) measurements, separate measurements were taken for the posterior and the anterior PSM, and were taken at the halfway point (AP) of each region. While the ML axis keeps the same orientation between posterior and anterior PSM, the DV axis does not, in that the tail is curled ventrally. The two separate height measurements take this ventral curl into account. Somite measurements for each axis were also taken through the middle of the tissue.

#### Cell tracking

Automatic nuclear tracking in Imaris involves two steps: spot creation (as described above) and spot tracking. Spot tracking requires an algorithm input, for which there are five options in Imaris: Connected Components (CC; Brownian Motion (BM); Autoregression Motion (AM); Autoregressive Motion Expert (AME); and Lineage (LI).

To determine which was the best tracking algorithm, we tested all five on the first hour of a two-photon movie (M3, Movie S3). For a first round of validation, tracks of the whole tailbud were made using the same spot reconstruction parameters, but different spot tracking algorithms. For each algorithm (except CC which does not include the parameter), two different Max Distance parameters were tested: 5 μm and 10 μm. Gap Size was set to 3 for all three AM-based algorithms. Various Intensity Weight parameters (ranging from 0 to 10,000) were tried for AME. In this first round of testing, the main result being assessed was track duration - any algorithm which could not track cells for long durations (> 30 mins) were to be excluded. The results of track duration are shown in Figure S4A. Regardless of parameters, CC and AME failed to track cells for long durations, and so these were excluded.

To determine which was the best tracking algorithm out of the remaining three (AM, BM, and LI), we manually validated the accuracy of PSM tracks. First, we filtered tracks by duration to only include those lasting the full hour. Then we manually selected 20 PSM tracks from each set of tracks. Manual validation involved using the same spot validation method as previously described, to follow the coloured spot over time and check if it stayed in the same nucleus. The initial LI track set (parameters: MD = 5, GS = 3) reported 8 cell divisions; however, manual validation confirmed these were all false positives. Given this level of inaccuracy, LI was excluded from further testing.

To choose between the two remaining algorithms (AM and BM), three different track sets were made for each algorithm, one for each MD parameter: 5, 7.5, and 10. Gap Size was kept constant (GS = 3) for all track sets. The results of validation are shown in Figure S4B. The best tracking algorithm and parameter set for this movie (M3) was found to be Autoregressive Motion (MD = 5), with 16/20 tracks showing no errors. Higher MD values led to more errors for both algorithms, and more gaps for AM. Track gaps were assumed to be a possible source of error, so it was noted whether errors occurred over gaps. However, for all track sets, gaps showed mostly accurate tracking, and for AM (MD = 5), all three gaps were error-free.

After choosing Autoregressive Motion as the best algorithm, this was tested on the other movies (M1, M2), using different MD parameters, to check if MD needed adjusting for differences in frame interval. For both M1 and M2 two sets of AM tracks were made, one with MD = 5, and one with an “adjusted” MD (MD = 3 for M1, MD = 6 for M2). For each set of tracks, track durations were measured, and PSM tracks were manually validated as before Figure S4C.

The results show that M1 (AM, MD = 5) tracks were highly accurate (20/20 accurate PSM tracks) but, on average, did not last very long (mean = 25 min). Adjusted MD tracks were not validated for M1, as they showed even shorter average duration (mean = 10 min). M2 showed highly inaccurate tracks for both MD values, but less so for MD = 6 (10/20 accurate PSM tracks). The difference in accuracy between M1 and M2 is likely due to the difference in frame interval (70s for M1, 180s for M2). The short track durations for M1 is likely due to slight twitching of the embryo during imaging, causing track breaking.

Based on all these results, Autoregressive Motion (AM) was used as the tracking algorithm for all three movies, with Gap Size set to 3 for all movies. For M1 and M3, a Max Distance (MD) of 5 was used, but for M2 an MD of 6 was used. Because M3 showed both long track durations (mean = 50 min) and high tracking accuracy, this movie was used as the main movie in tracking analyses, with the other two movies used as additional movies to check the generality of results.

## 3. Results

### 3.1. The zebrafish presomitic mesoderm decreases in volume as it extends

To better understand the tissue dynamics involved in zebrafish paraxial mesoderm elongation, a long-term, three-dimensional approach is required to determine the overall shape changes associated with elongation. We first made use of *in situ* hybridization chain reaction (HCR) to stain for the PSM markers *msgn1* and *tbx6* in embryos fixed at various stages of somitogenesis, together with DAPI to mark nuclei (Figure 1A). We focused on embryos past the 16 somite-stage to ensure that direct convergence of cells into the paraxial mesoderm from more lateral regions was no longer happening. Although the exact stage at which this process stops is not known, previous work has shown that cells can directly enter the somitic mesoderm of up the 17^th^ somite, without having derived from the tailbud (Steventon et al., 2016). 3D reconstructions of the PSM were created by generating 2D contours around the tissue, at regular z-intervals, for each image (Figure 1B; Figure S1A-C; Movie S1). To obtain information about cell numbers over time, we created “spots” automatically using Imaris. The nuclei (DAPI signal) from each PSM/somite surface were isolated (Figure 1C) and spots were generated from these nuclei (Figure 1D). Together, these measurements provided information about tissue lengths, volumes, and cell numbers over time.

Measurements were taken of the PSM and the nascent somite at each stage. This allowed us to measure changes to the PSM over time and the size of each nascent somite at its time of formation. It also enabled us to calculate length, volume, and cell number values for the whole paraxial mesoderm (from the 16^th^ somite onwards) in a way that eliminated any growth or compaction happening in the somites after their formation. In other words, we calculated the length, cell number, and tissue volume of the 16 somite-stage PSM from this stage until the end of somitogenesis, including the tissues it gives rise to, but excluding processes not relevant to PSM elongation.

Measurements of the zebrafish PSM length (Figure 1E), cell number (Figure 1F) and volume (Figure 1G) all showed depletion through time, as somites are continually being generated at a faster rate during these stages than any expansion or elongation of the PSM.. Additionally, each nascent somite is smaller than the previous one, and so the nascent somites display a similar gradual reduction in length (Figure 1H), cell number (Figure 1I) and volume (Figure 1J) as observed for the PSM. These values then enabled us to calculate how the paraxial mesoderm elongates over-time, by summing values for each nascent somite and the PSM at each somite stage. Rather than using specific measurements, trendline equations were calculated from the data - in order to obtain average values for each stage that had more than one data point. The following function *f*(*x*) can then be expressed in terms of somite stage *x*, the PSM function *p*(*x*), and the nascent somite function *s*(*n*), which sums from *n* =17 to the given somite stage *x*. The equation, which can be applied to length, cell number, and tissue volume, is as follows:

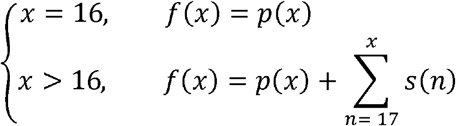

The results show that the length of the paraxial mesoderm increases considerably (~ 100%) throughout these stages of somitogenesis (Figure 1K; purple line). Taken together with the PSM measurements, this result shows that the PSM is elongating, but the length of each nascent somite specified is greater than the amount of elongation between somite stages, and so the length shows a gradual net decrease. Paraxial mesoderm cell number initially stays constant (when elongation is fastest) before showing a very slight increase (~ 10%) at later stages (Figure 1L; purple line). This late, slight increase in cell number is likely due, in part, to NMp addition - given that NMps only contribute to the paraxial mesoderm at these same late stages (Attardi et al., 2018). Again, taken with the PSM measurements, this shows that the PSM does show an increase in cell number, but the number of cells lost to the somites is greater than this increase between somite stages, and so the cell number shows a net decrease. Interestingly, paraxial mesoderm tissue volumes do not show the same trends as those of cell numbers (Figure 1M; purple line). While both PSM and nascent somite volumes decrease exponentially (like cell numbers), this decrease is steeper. As a result, the paraxial mesoderm volume does not remain constant with a late increase, but instead decreases.

Taken together, these results show that the paraxial mesoderm elongates considerably over time. However, this elongation is not the result of growth. While the number of cells does increase slightly, this does not cause tissue expansion – as the tissue decreases in volume over time. Therefore, the paraxial mesoderm elongation is coupled with an overall volume decrease in zebrafish embryos.

### 3.2. Compression is observed across both dorsal-ventral and medio-lateral axes

To determine in which axes the tissue is deforming to generate this increase in length, measurements of the both the posterior and the anterior PSM were taken along both the dorsal-ventral (DV; Figure 2A; Movie S1) and medio-lateral axes (ML; Figure 2B; Movie S1). These showed that both height and width decrease over time by ~ 50% (Figure 2C). To determine whether anterior PSM thinning is actively occurring in that region of the tissue or the result of being progressively generated from a thinning posterior PSM, we used photolabelling and live imaging to follow how groups of cells deform over time. Embryos were injected at the one-cell-stage with mRNA for KikGR – a fluorescent protein that switches from green fluorescence to red fluorescence upon photoconversion with UV light (Habuchi et al., 2008). Dorsoventral stripes (across the full height of the PSM) were labelled in both the progenitor region and anterior PSM (Figure 2D,E; Movie S2). The length (AP axis) and height (DV axis) of these labels were measured immediately after labelling, and again after 2 hours. By accumulating measurements from labels placed at different regions across the PSM, a clear trend is observed with increased convergence and extension observed in the most posterior region. The length-height ratio triples in posterior-most labels but stays constant in anterior-most labels (Figure 2F). Given that anterior PSM labels show very little deformation, this suggests that anterior PSM thinning is mostly the result of this tissue being progressively generated by a thinner posterior PSM at each preceding stage and supports the notion that the active region of deformation is within the posterior progenitor domain.

**Figure 2:**
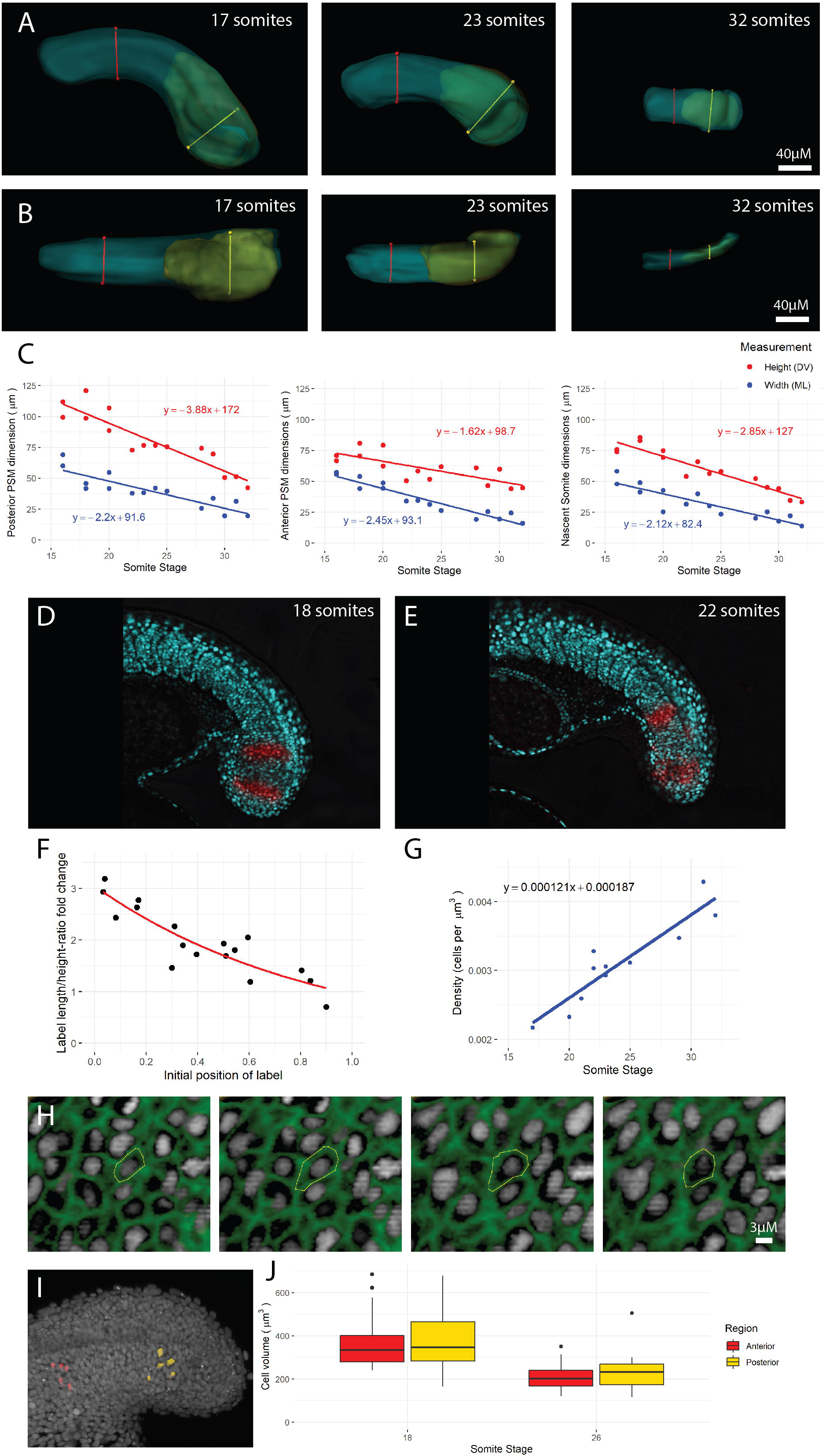
The PSM is compressed in height and width over time. Height (DV axis) (A) and width (ML axis) (B) measurements were taken of the posterior PSM (yellow line), anterior PSM (red line), and nascent somite (not shown) from the 16 somite-stage to the end of somitogenesis. (C) All height and width measurements show a decrease over the course of somitogenesis. (D, E) Using a photoconvertible protein, dorsoventral stripes of cells in the PSM were photolabelled (red) and imaged over four somite-stages (n = 7 embryos). (F) The height and length of each label at the beginning and end of imaging was measured, along with the initial distance of the label from the posterior end of the PSM. These measurements were used to calculate a length/height ratio fold change over time, which is plotted over initial position AP position of the label (normalized from 0 to 1 between embryos, with 0 being the posterior end and 1 being the nascent somite posterior boundary). (G) PSM density (cell number divided by tissue volume) increases over the course of somitogenesis. (H) Phalloidin-stained (green) tails of 18 somite-stage and 26 somite-stage embryos were used to measure cell volumes, by drawing contours around the cell membrane at every z-slice. This was done for 5 randomly selected posterior cells and 5 randomly selected anterior cells per embryo (n = 10 embryos). (I) 3D surfaces of posterior (yellow) and anterior (red) PSM cells. (J) Boxplot showing cell volumes between regions and between stages. Cell volumes are not significantly different between regions (t_(78)_ = −0.36, p = 0.72) but are significantly different between stages (t_(78)_ = 6.39, p < 0.001).

The result that the paraxial mesoderm shrinks in volume over time suggests that PSM density increases over time, pointing to a mechanism by which the PSM progenitor region compresses in DV and ML axes, to extend along the AP axis. This was found to be the case, by dividing the number of PSM cells by the PSM tissue volume to obtain a density measurement in terms of cells per μm^3^ (Figure 2G). This density increase is substantial (~ 80%), which suggests that cells must themselves decrease in volume to account for tissue-level compression. To test this, we fixed two batches of embryos (one at the 18 somite-stage, and one at the 26-somite stage, n = 5 embryos per batch) and stained the cell membranes with phalloidin to segment out individual cell shapes (Figure 2H) in both the posterior (Figure 2I; yellow) and anterior (Figure 2J; red) PSM. The results show that within each stage, there was no difference between anterior cell size and posterior cell size (Figure 2J; t_(78)_ = −0.36, p = 0.72). However, there was a significant difference between stages (Figure 2J; t_(78)_ = 6.39, p < 0.001) - PSM cells of 26 somite-stage embryos (mean = 220 μm^3^) were smaller than those of 18 somite-stage embryos (mean = 367 μm^3^). Importantly, while cell sizes could be reducing due to cell divisions without subsequent cell growth, the number of cells in the paraxial mesoderm remains relatively constant over time (Figure 1L, purple line). Therefore we can rule out the possibility that cell volume decrease over time is the result of cell division, and confirm that it is an active process of compression. Together, these results demonstrate that the PSM progenitor domain undergoes a compression-extension deformation that is progressively transmitted to the anterior PSM as cells displace from posterior to anterior.

### 3.3. Generation of 3D cell tracking data

To determine the cell behaviours and movements underlying the tissue-level compression extension of the zebrafish PSM, individual cell 3D tracks are required. To obtain cell tracks, we live-imaged zebrafish embryos spanning the 14-26 somite stages (Figure S2). To draw meaningful conclusions from cell tracks, it is important to measure cell displacement with respect to appropriate frames of reference. The zebrafish tailbud is elongating and uncurling throughout most of somitogenesis, and so tracking PSM cells through absolute space will simply reflect the global tissue movement. Using Imaris imaging analysis software, a “reference frame” was placed at the end of the tailbud, at the DV and ML midline (Figure S3). The reference frame includes axes, which were orientated to match the three biological axes (x = AP, y = DV, z = ML) and adjusted every 5 frames for the full movie duration, for each movie. Imaris then automatically calculated the movement and rotation of the reference frame, using linear interpolation, for in-between frames, and placed the reference frame appropriately for these frames. This allowed visualisation (of movies/tracks) to reflect the normalisation for global movement.

Having chosen the best tracking algorithm and tracking parameters (see methods and Figure S4), we manually selected paraxial mesoderm tracks (Figure 3A,C yellow spots) from all cell tracks in the dataset (Figure 3A,C; grey spots). As in surface reconstructions (Figure 1), only one lateral half of the paraxial mesoderm was included – cells past the midline of the embryo were excluded. Because the images only contained nuclear signal, and no gene expression information (unlike the HCR images used for surface/spots reconstruction (Figure 3B, D; cyan and magenta spots)), it is likely that some notochord and neural progenitors are included in the tracks. However, these will constitute a negligible minority of cells, especially as any tracks that moved into the notochord or neural tube were excluded. The resulting cell tracks are shown colour-coded by time (Figure 3E; Movie S3).

**Figure 3:**
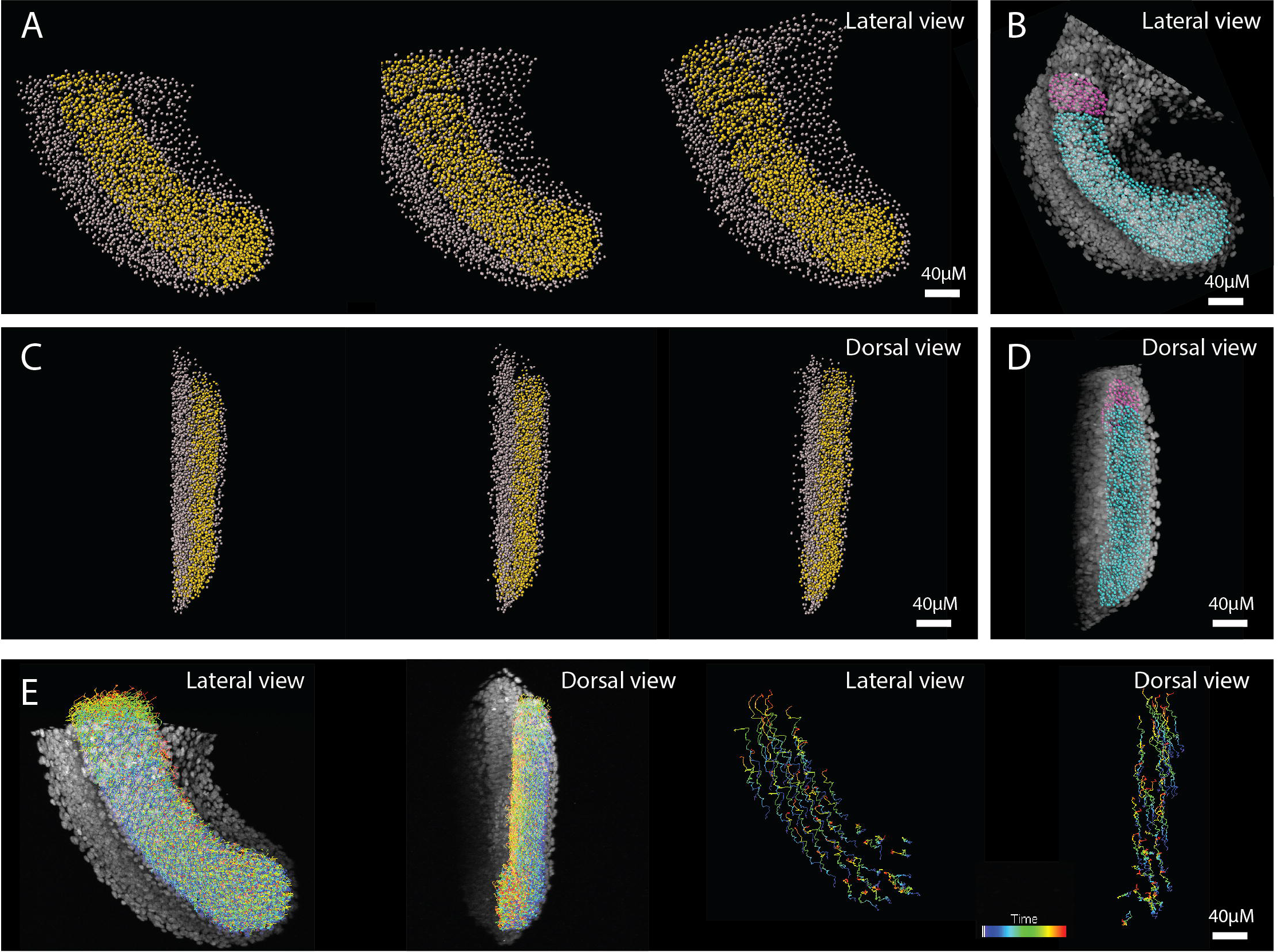
Paraxial mesoderm cell tracks. Tracks were generated of the whole tailbud and all non-paraxial mesoderm tracks were then manually removed. Paraxial mesoderm (PM) tracks (yellow spots) and non-paraxial mesoderm tracks (grey spots) shown at 0, 1, and 2 hours after imaging. (B) PM spots reconstructions (cyan: PSM & pink: nascent somite) from a similar-stage HCR image is shown for comparison/validation of selection accuracy. (C, D) The same images as above, but from a dorsal view: anterior is top, medial is left. (E) All PM tracks of full movie (colour-coded by time) superimposed over first frame image, shown for lateral and dorsal views (left images). A random selection (n = 50) of long-duration (>2 hr) PM tracks are shown, for lateral and dorsal views (right images).

### 3.4. PSM elongation is associated with non-directional cell rearrangement

We first set out to compare how cells alter in the rates of cell rearrangement as they transition from the posterior (Figure 4A; shaded red to yellow) to anterior PSM (Figure 4A; blue). To analyse this in 3D, we performed a nearest neighbour analysis that takes a given number *k* of nearest cells (neighbours) for each cell to form a neighbourhood for that cell at every frame. We then calculate the number of new cells that enter this neighbourhood over a given time *t* (Figure 4A).

**Figure 4:**
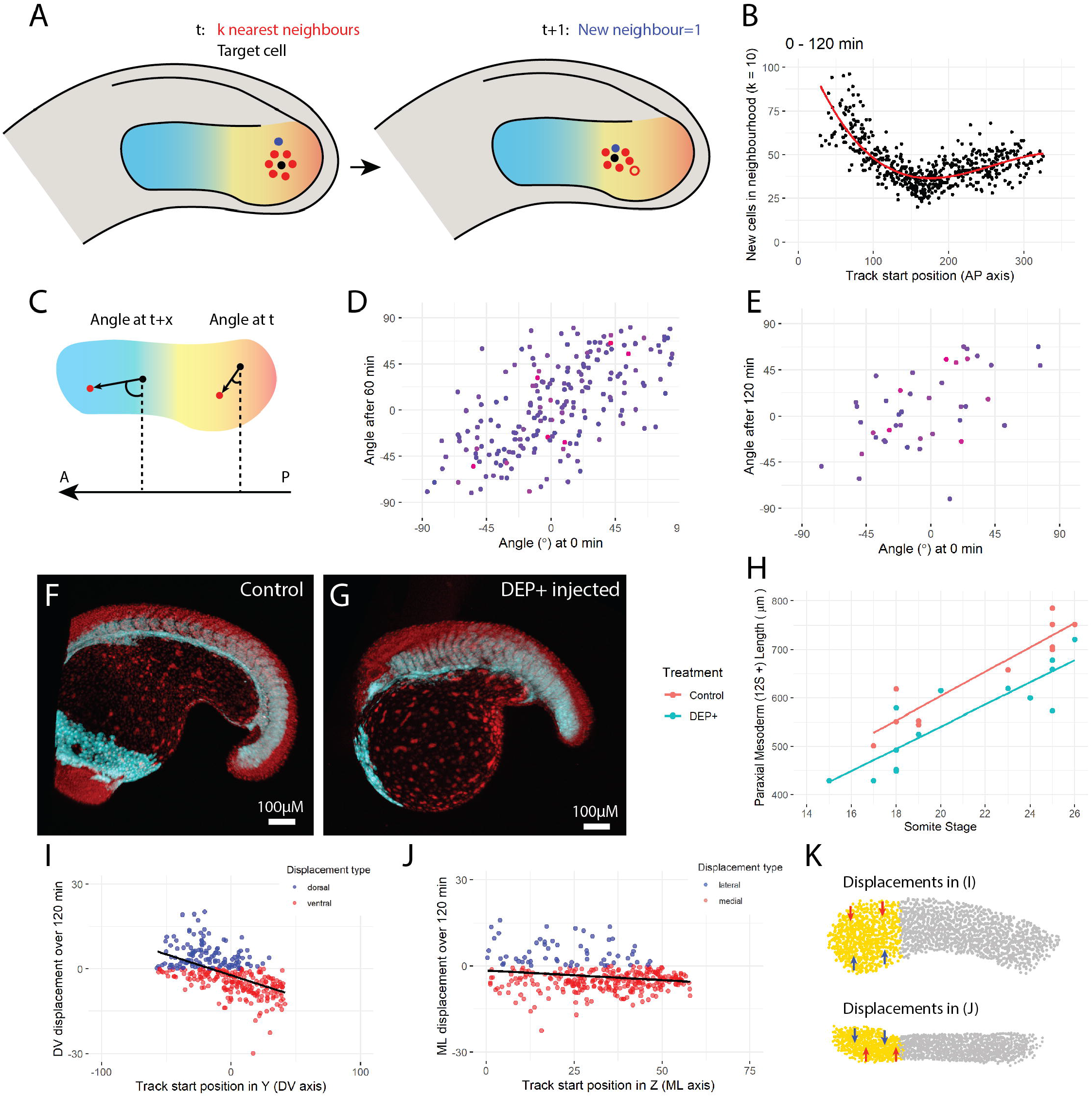
Posterior PSM cells drive convergent extension, but not via directional intercalation. (A) Analysis of neighbour exchanges. For each cell (black), a neighbourhood was specified as a given number (k) of nearest neighbours (red cells). After a given time interval, the number of new cells (blue) that entered the target cell’s neighbourhood was calculated. In this example, one new cell joins the neighbourhood (k = 6) over time, with the cell that is no longer in the neighbourhood shown in red outline. (B) The number of new cells entering each target cell’s neighbourhood (k = 10) over 120 min is plotted against the initial AP position of the target cell, with 0 being the posterior end of the tail, and ~ 300 being the anterior-most cells. The results show more cell mixing in the posterior than the anterior PSM (C) Analysis of neighbour angle changes. For each cell (black), the angle between the vector to the nearest neighbour (red) and the AP axis was calculated at the initial timepoint (t). After a given time interval (t + x), the same vector angle was calculated between the two cells (regardless of whether they were still neighbours). An angle of 0° means the cells lie perpendicular to the AP axis, while an angle of 90° or −90° means the cells lie parallel to the AP axis. Measured neighbour angle changes are plotted after 60 min (D) and 120 min (E), with the later angle plotted against the initial angle. Points are colour coded by the change in distance between the neighbours (blue to red: low to high). Neither graph shows evidence of neighbours lining up along the AP axis (angles of 90°/-90°). *tbx16::GFP* (cyan) x *h2a*::mCherry (red) embryos were either left uninjected (control, n = 6) (F) or injected with DEP+ (n = 7) (G) and then live-imaged. (H) The length of the paraxial mesoderm (from the 12th somite onwards) was directly measured for these embryos over time. (I) Strong DV axis convergence over 120 min: dorsal cells (x > 0) move ventrally (y < 0, red) and ventral cells (x < 0) move dorsally (y > 0, blue). (J) Weak ML axis convergence over 120 min: lateral cells (high x) move medially (y < 0, red) and medial cells (low x) move laterally (y > 0, blue). (K) Schematic summarising displacements of posterior PSM cells. Top view is lateral, showing DV convergence. Bottom view is dorsal, showing ML convergence.

The results of this analysis show that posterior cells exchange neighbours more than anterior cells (Figure 4B) that is consistent with measurements of mechanical properties of the tissue (Mongera et al., 2018). These results were consistent between small (*k* = 10) and large (*k* = 50) neighbourhoods, and between movies (Figure S5A). Given that an artificial increase in new neighbours might be caused in a region where cells are on average tracked for shorter duration (i.e. if average track duration was much lower in the posterior then new cell tracks might be interpreted as new neighbours), it was important to check that the observed trends were not simply an artefact of a possible correlation between track duration over track start position. However, there was no clear correlation between track duration and track start position (Figure S5B).

To qualitatively explore the directionality of cell movements within the PSM, we next measured the AP displacement of tracks over time. We used three reference frames along the paraxial mesoderm: the tailbud tip, the posterior end of the notochord proper, and the posterior boundary of the start-of-movie nascent somite (Figure S6). The results show that, relative to the tailbud tip or the posterior notochord, almost all cells displace anteriorly (Figure S6A,B). However, relative to the nascent somite, almost all cells displace posteriorly (Figure S6C). These results highlight the difficult of defining movement as anterior vs posterior in an elongating tissue. While it is certainly the case that cells disperse along the AP axis, defining this dispersal as anterior vs posterior is purely subjective, based on which end of the embryo displacement is measured relative to. Given that the anterior end of the tissue/embryo is equally valid as a reference point as the posterior end is, there is no reason to define the AP dispersal as anterior movement. Therefore, we set out to determine the angular changes in cell displacement within the progenitor domain.

To determine the directionality of cell rearrangement we measured angle changes between neighbours, relative to the axis of elongation, over time. This involves taking the nearest neighbour for each cell at the start of the movie, taking the vector between those two cells, and the angle between this vector and the AP axis. Then, after a specified time-period *t*, the same two cells are taken (regardless of whether they are still neighbours), and their vector angle relative to the AP axis is measured again (Figure 4C). In this way, the orientation of cell pairs along the AP axis can be measured over time, to test if neighbours are aligning with each other along the axis to drive elongation. In a situation where cells are directionally rearranging to drive convergence and extension along the AP axis, it would be expected, irrespective of the initial angle of nearest neighbours, cell pairs should converge towards either 90° or −90° at t+1 (i.e. the top and bottom portions of Figure 4D,E). Interestingly, while there is substantial rearrangement happening between neighbour pairs, this rearrangement does not suggest directional intercalation. Neighbour pairs are not orientating along the AP axis (Figure 4D), even when this is measured over long time periods (of 2 hours; Figure 4E). These results show that neighbour rearrangements are randomly orientated within the progenitor domain.

During zebrafish gastrulation, cells converge across the yolk and extend to form the tailbud-stage embryo. This convergent extension is dependent on the Planar Cell Polarity (PCP) pathway (Roszko et al., 2009). To experimentally rule out the possibility of PCP-driven convergence and extension behaviours during posterior body elongation, we injected mRNA for a dominant-negative form of *Xenopus* Dishevelled (DEP+) into one-cell-stage *tbx16::GFP* embryos (to mark the paraxial mesoderm). This form of Dishevelled can still function normally in canonical Wnt signalling, but not in the PCP pathway, due to deletion of the DEP domain (Tada and Smith, 2000). Injections resulted in post-gastrulation deformities with somites 1-12 showing a clear reduction in AP length (Figure 4G) as compared to uninjected controls (Figure 4F). However, despite the post-gastrulation paraxial mesoderm being shorter between injected and uninjected embryos, the elongation rate was similar (Figure 4H). This suggests that, while the convergent extension movements involved in paraxial mesoderm *formation* (during gastrulation) were perturbed, those driving paraxial mesoderm *elongation* were not affected. These results show that convergent extension movements during paraxial mesoderm elongation are not simply a continuation of earlier gastrulation movements, in terms of both an absence of orientated cell rearrangement and PCP pathway dependence.

As morphometric data showed convergence to the midline at the tissue level (Figure 2C) and photolabels confirmed this for the DV axis (Figure 2F), we next measured track displacements in the DV and ML axes in the progenitor region. All tracks that began, in the first frame, between the tailbud tip and the notochord proper were selected for analysis. For both DV and ML analyses, the tailbud tip reference frame was used, as this was the best reference frame to give a constant, correct midline for this region. The results show that posterior PSM cells do converge in the dorsoventral axis, with ventral cells displacing dorsally and dorsal cells displacing ventrally (Figure 4I). ML displacement was also observed, although to a lesser degree (Figure 4J) – as would be expected from previous tissue measurements showing a greater absolute decrease in height than in width. These displacements are summarised in Figure 4K.

These results further confirm that the PSM is undergoing compression-extension, as PSM cells displace towards the midline of the tissue (in both DV and ML) while dispersing along the AP axis.

## 4. Conclusions and Discussion

We propose that the zebrafish PSM elongates via a compression-extension mechanism based on the following observations. Firstly, elongation is associated with a decrease in tissue volume, with a concomitant increase in cell density and a decrease in cell volumes. Secondly, cells converge to the midline in both DV and ML axes, while dispersing along the AP axis. Finally, elongation occurs in the absence of directional cell rearrangement and in a PCP-independent manner. The latter observation rules out an alternative hypothesis of classical convergence and extension cell behaviours known to be important for the elongation of the embryonic axis during gastrulation (Roszko et al., 2009). An absence of directional intercalation between neighbours has been observed in zebrafish through following individual transitions in boundary contacts over short time-scales (Mongera et al., 2018). In addition by tracking cells and subtracting their movements from the average movements of neighbouring cells, it has been previously shown that the posterior PSM displays a disordered cell movement (Lawton et al. (2013)). Here, we confirm and extend these findings by showing that even after 2 hours, cells do not show any evidence of orientating themselves along the AP axis. Taken together, these results support a model in which non-directed cell movements are coupled with DV and ML convergence as the PSM compresses and extends along the AP axis.

That PSM elongation in zebrafish is associated with non-directional cell rearrangements reflects closely what has been observed in avian embryos (Bénazéraf et al., 2010). However, in the random motility model of avian PSM elongation, an important component is a density gradient along the AP axis. In this model, a low density of cells in the posterior (versus a high density of cells in the anterior) ensures that cell addition from the node causes the PSM to expand posteriorly. In zebrafish, there is no AP density gradient - instead, the posterior PSM appears to be more fluid-like due to decreased cell-cell adhesion (Mongera et al., 2018). This suggests that while an AP gradient in PSM tissue fluidity is conserved across vertebrates, the mechanism responsible for this is different among species (cell density in chick vs cell-cell adhesion in zebrafish). Alternatively, it could be that compression-extension is a feature of tailbud stages of somitogenesis. Further studies in avian embryos measuring cell density over longer time periods – and including tailbud stages – would help answer these questions.

Although we show that the increase in PSM density is associated with a decrease in cell volumes, we do not know the causal relationship between these processes. Particularly, whether the mechanism is internal (i.e. the PSM is compressed from within due to cells contracting) or external (i.e. the PSM is compressed from outside, causing cells to shrink). The latter seems more likely, given the high levels of cell mixing and displacement. If the PSM was being externally compressed (either by outer PSM cells, or by other tissues), this would likely lead to a mixture of high cell displacement, undirected neighbour rearrangements, and cell shrinkage – all of which are happening in the zebrafish PSM. A likely mediator of the balance between tissue intrinsic cell movements and extrinsic forces is the extracellular matrix (ECM). Both PSM elongation and somitogenesis are dependent on integrins and fibronectin in zebrafish embryos (Dray et al., 2013; Jülich et al., 2009), and appropriate remodelling of the fibronectin matrix is mediated by the cell adhesion molecule Cadherin2 during somite formation (Jülich et al., 2015). How such mechanisms operate at external boundaries to the PSM over time, and as cells transit through the tissue will likely reveal new insights into the mechanisms of tissue elongation.

In addition to tissue-ECM interactions, it is likely that forces will be acting between adjacent tissues to help coordinate multi-tissue morphogenesis during body axis formation. Indeed, frictional forces acting between the notochord and PSM have been shown to be important for shaping the chevron-like shape of somites as they mature (Tlili et al., 2019), and anterior notochord expansion due to vacuolation is required for the continued elongation of the somatic compartment in already segmented regions of the body axis (Mclaren and Steventon, 2021). How such multitissue interactions operate within the tailbud of amniote embryos has been recently addressed in a study that demonstrates a key role for PSM expansion in timing the entry of new cells to the posterior progenitor domain (Xiong et al., 2020). Taken together, these studies reveal posterior body axis elongation to be a tractable model system to better understand the role of multi-tissue mechanical interactions in morphogenesis. This work further enables this by providing a quantitative understanding of the tissue shape changes and cell rearrangements associated with PSM compression-extension.

## Supplementary figure legends

**Figure S1: Tissue reconstructions of the PSM and somites.**

(A) 2D contours (yellow outlines) manually drawn around the PSM at regular z-slices, up to the midline. (B) Maximum projections showing all PSM contours, from lateral, dorsal, and posterior views (left to right). (C) Same images as in B but showing PSM surface (cyan). (D) Contours and surface (pink) of the nascent somite (left, middle), and automatic *msgn1* surface (gold, right). All images are of the same 23 somite-stage embryo. Scale bar: 50 μm.

**Figure S2: Two-photon time-lapse imaging of the zebrafish tailbud.**

Three embryos of different stages were live-imaged for 2-3 hours each. Nuclei are labelled with either *h2a*::mCherry or *h2b*::GFP (both shown as grey). Images are shown of the tailbud at 0, 1, and 2 hours after the start of imaging. All images are maximum projections. Scale bars: 40 μm. Each movie is summarized in the table below, in terms of somite stages, frame interval, and nuclear label.

**Figure S3: Normalizing cell movements for global tail uncurling.**

A “reference frame” was placed at the DV/ML midline of the tailbud tip (with axes lined up to match biological axes) at regular intervals. (A) Individual z-slice showing reference frame placement over 2 hrs. Scale bar: 50 μm. (B) Images and cell tracks before (left) and after (right) “resampling” images to fix the reference point (tailbud tip) in space. Images and tracks (all cells) show a large dorsal movement before resampling, but none after resampling. Tracking colour code (blue to red) indicates time. Scale bars: 40 μm. All images from M3. Anterior is top, dorsal is left.

**Figure S4: Validation of automatic tracks.**

(A) Tracks were generated of the whole tailbud (first hour of movie M3) using 5 different algorithms: Autoregressive Motion (AM); Autoregressive Motion Expert (AME); Brownian Motion (BM); Connected Components (CC); and Lineage (LI). For each algorithm, two sets of tracks were made for two Max Distance (MD) parameters (5, 10). Boxplots show track duration for each set of tracks. AME and CC were excluded from further analysis due to short track durations. (B) Manual validation of track accuracy (LI was excluded at the beginning of analysis due to high false positive reporting of cell divisions). Tracks were generated for AM and BM, for three MD parameters each (5, 7.5, 10), and 20 PSM tracks (with duration > 60 min) were randomly selected for validation over 60 min. Number of errors (top left) and number of accurate tracks (error-free tracks) (top right) were recorded. Track gaps were also recorded (bottom left), and the percentage of accurate (error-free) gaps were noted (bottom right). (C) Having identified AM as the best algorithm, and MD = 5 as the best value for movie M3, tracks were made for all three full movies. For M1 and M2, two sets of tracks were made with an adjusted (for difference in frame interval) and non-adjusted MD value. Boxplot shows track duration for each set of tracks. For each track set, 20 PSM cells were randomly selected and validated as before. Bar charts show number of errors (left) and number of accurate tracks (right). (For M1, the adjusted MD tracks were not validated, given the perfect accuracy and longer duration of non-adjusted MD tracks.)

**Figure S5: Neighbourhood analysis.**

Analysis was performed using MATLAB scripts written by L.M. (A) Neighbourhood analysis results for all three movies (left to right), over a time period of 120 min, for small (k = 10, top row) and large (k = 50, bottom row) neighbourhoods (note different y-axis scales). Each point represents the neighbourhood of one cell. The number of new cells which entered that cell’s neighbourhood is plotted against the starting position (AP) of that cell (0 = posterior). The results show a strong gradient of cell mixing, from the posterior (high mixing) to the middle (low mixing) of the paraxial mesoderm. From the middle to the anterior, there is a slight increase, likely related to somite morphogenesis. These results are highly similar to those of photolabelling experiments, confirming the reliability of cell tracks. (B) Track duration plotted against track start position, for each movie, to confirm that these patterns are not an artefact of track durations.

**Figure S6: Multiple biological reference frames (RFs) along the AP axis.**

(A) Individual z-slice showing tailbud (TB) RF as previously described (Figure 17); notochord (NC) RF placed at the posterior end of the notochord proper (where the notochord “funnels out” into the notochord progenitors); and somite (SOM) RF placed at the posterior boundary of the start-of-movie nascent somite. Time is shown from left to right. (B) All three RFs shown together on maximum projection images from lateral (top) and dorsal (bottom) views. Scale bars: 40 μm. All images from two-photon movie M3. Anterior is top, dorsal is left (except bottom row in B, in which medial is left).

**Figure S7: Anteroposterior displacement is relative to the reference frame.**

Measurements of AP displacement of each track over 120 min are shown for all three movies (top to bottom), relative to three different reference frames: tailbud tip; notochord proper end; and the start-of-movie nascent somite (left to right). AP displacement is shown on the y-axis, with positive values indicating anterior displacement, and negative values indicating posterior displacement (points are also colour coded for displacement type to highlight this). This is plotted against track start position, where 0 is the reference frame position (negative values are posterior to this, positive values are anterior). The data shows that almost all cells move anteriorly relative to the tailbud tip, but also that almost all cells move posteriorly relative to the nascent somite, with displacement relative to the notochord showing similar results to that of the tailbud tip, but with slightly more posterior displacement. All trendlines (black) show a positive correlation i.e. anterior cells move anteriorly while posterior cells move posteriorly.

## Movie captions

**Movie S1: 3D morphometric measurement workflow.**

*in situ* hybridization chain reaction (HCR) for *msgn1* (yellow) and *tbx6* (red), with nuclei labelled by DAPI staining (grey). Image is a rotating 3D view of a confocal image. Each z-slice is then shown (scrolling from lateral to medial and back). Manual contours (cyan lines) were drawn around the PSM up to the embryo midline. A 3D reconstruction was generated from these contours (cyan surface), providing tissue volume information. DAPI signal was then isolated from this surface and used to generate a spot of each nucleus centre (cyan spheres), providing cell number information. Measurements of height and width of both the posterior (yellow lines) and anterior (red lines) PSM, as well as the length of the PSM (white line). The movie shows this process for the PSM at the 23 somite-stage, but the same process was done for the PSM, and the nascent somite, at each somite stage from the 16 to the 32 somite-stage.

**Movie S2: Convergent extension of photolabels.**

Zebrafish embryos were injected at the one cell stage with mRNA for KikGR, a photoconvertible protein which localizes to the nucleus of each cell (shown in cyan). Dorsoventral stripes (red) along the length of the PSM were photoconverted with a UV laser on a confocal microscope. The movie shows a live zebrafish tailbud with three labels along the PSM. The move shows 2 hours of imaging (at 28°C), which then rewinds to show original label positions again. The posterior label (bottom right) undergoes far more convergent extension than the middle and anterior labels.

**Movie S3: Paraxial mesoderm cell tracks.**

*h2b::GFP* zebrafish embryos were live imaged on a two-photon microscope at 28°C. A reference frame was placed at the end of the tail at regular time intervals, and images were then registered to this point, to eliminate tail movement/uncurling before 3D automatic tracks of all nuclei were generated. All cell tracks outside the paraxial mesoderm were manually removed, to create a tracking of only paraxial mesoderm cells. The movie shows imaging over 2 hours, with paraxial mesoderm tracks colour-coded by AP position. At each timepoint, paraxial mesoderm tracks shown movement over the previous 5 minutes of imaging.

## Acknowledgements

We would like to thank Berta Verd for discussions through the development of this work, and Toby Andrews for essential suggestions on image analysis. Thanks to the Cambridge Advanced Imaging Centre (CAIC) for imaging support. B.S. is supported by a Henry Dale Fellowship jointly funded by the Wellcome Trust and the Royal Society (109408/Z/15/Z). L.T. is supported by a scholarship from the BBSRC. L.M. is supported by an EPRC grant (EP/R025398/1).

