## Supplementary material for "The zebrafish presomitic mesoderm elongates through compression-extension": Figures S1-7

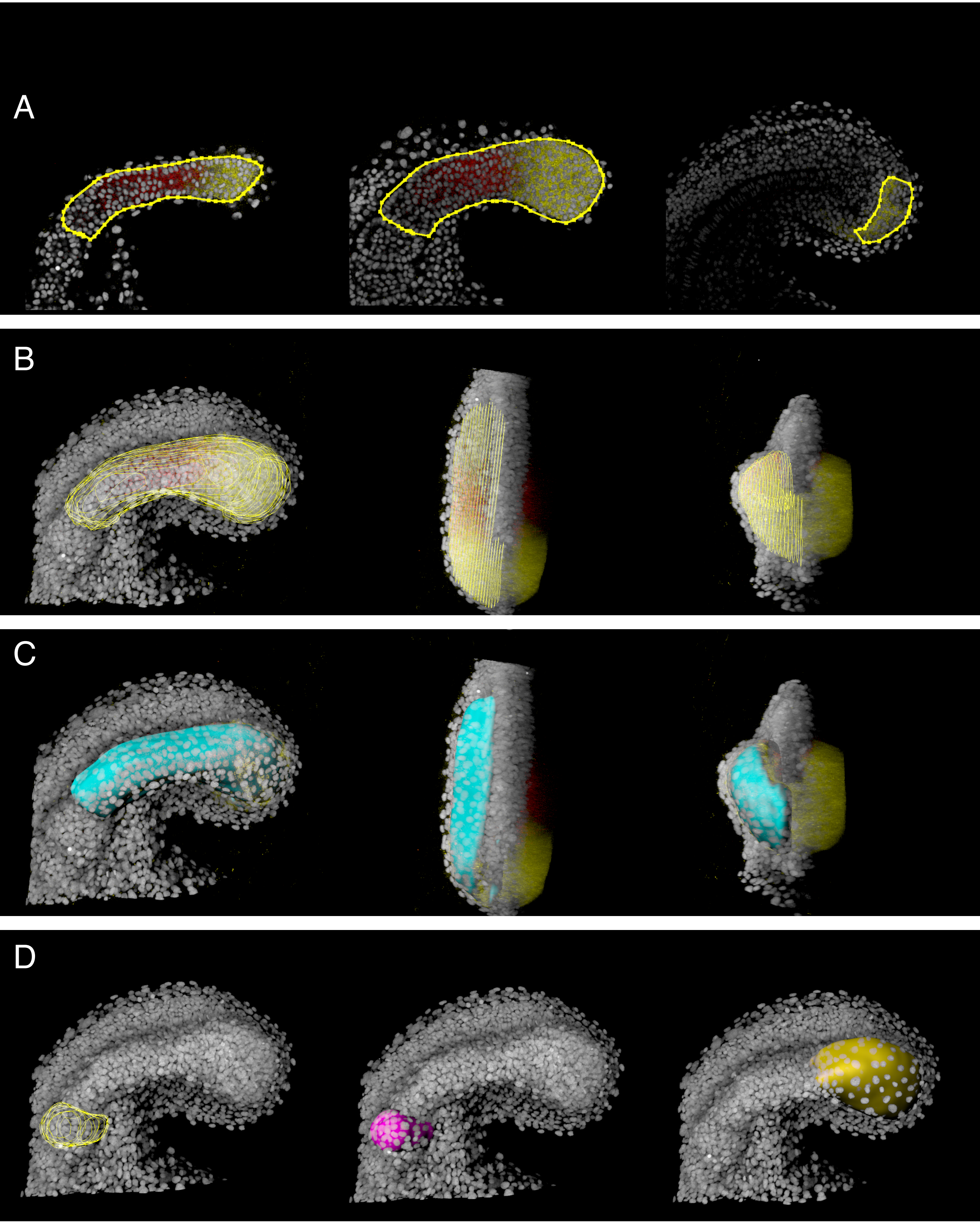


Figure S1

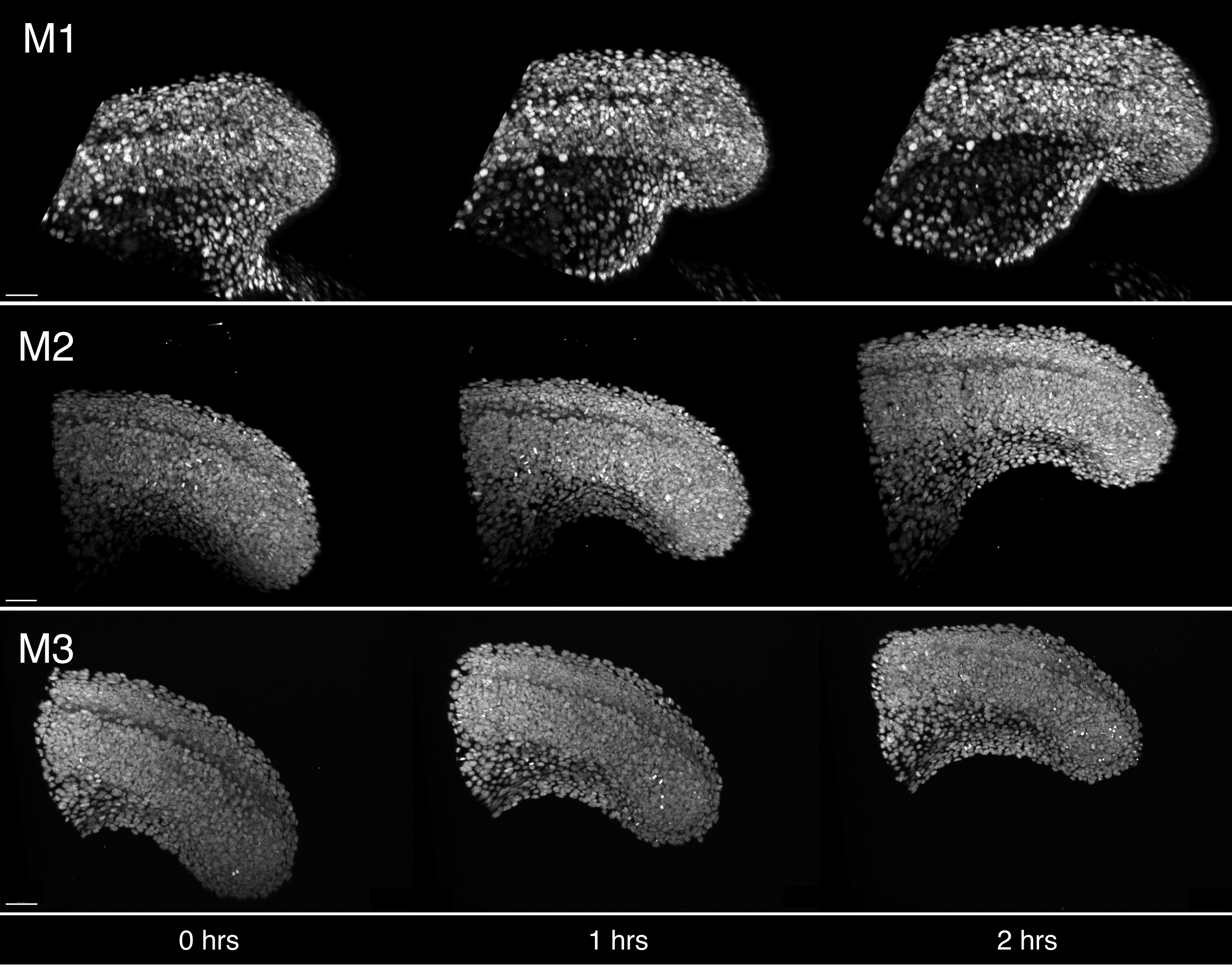


| Movie | Somite-stages | Frame interval (s) | Nuclear label |
|---|---|---|---|
| M1 | 14 - 20 | 70 | *h2a::mCherry* |
| M2 | 18 - 22 | 180 | *h2b::GFP* |
| M3 | 22 - 26 | 150 | *h2b::GFP* |

Figure S2

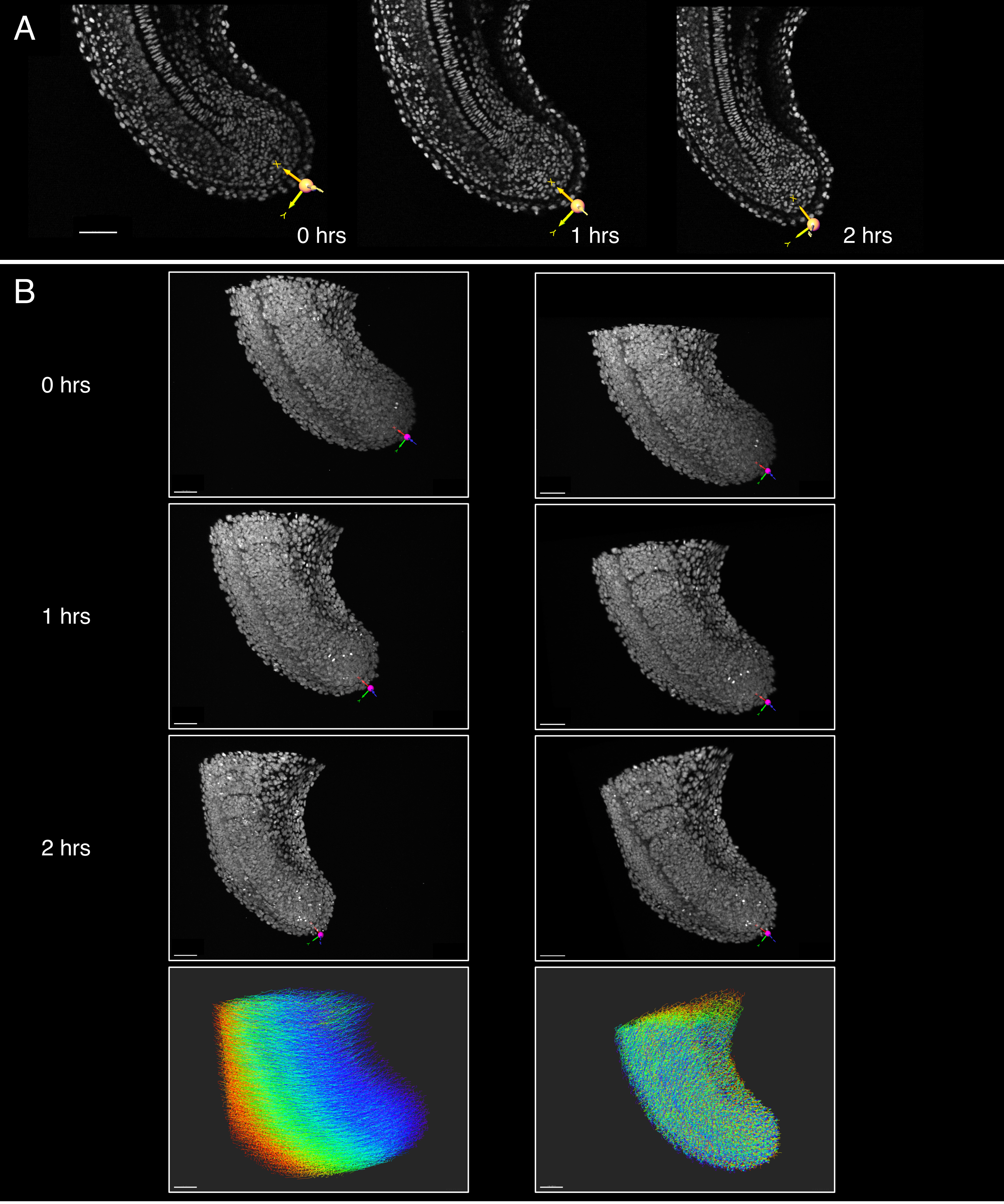


Figure S3

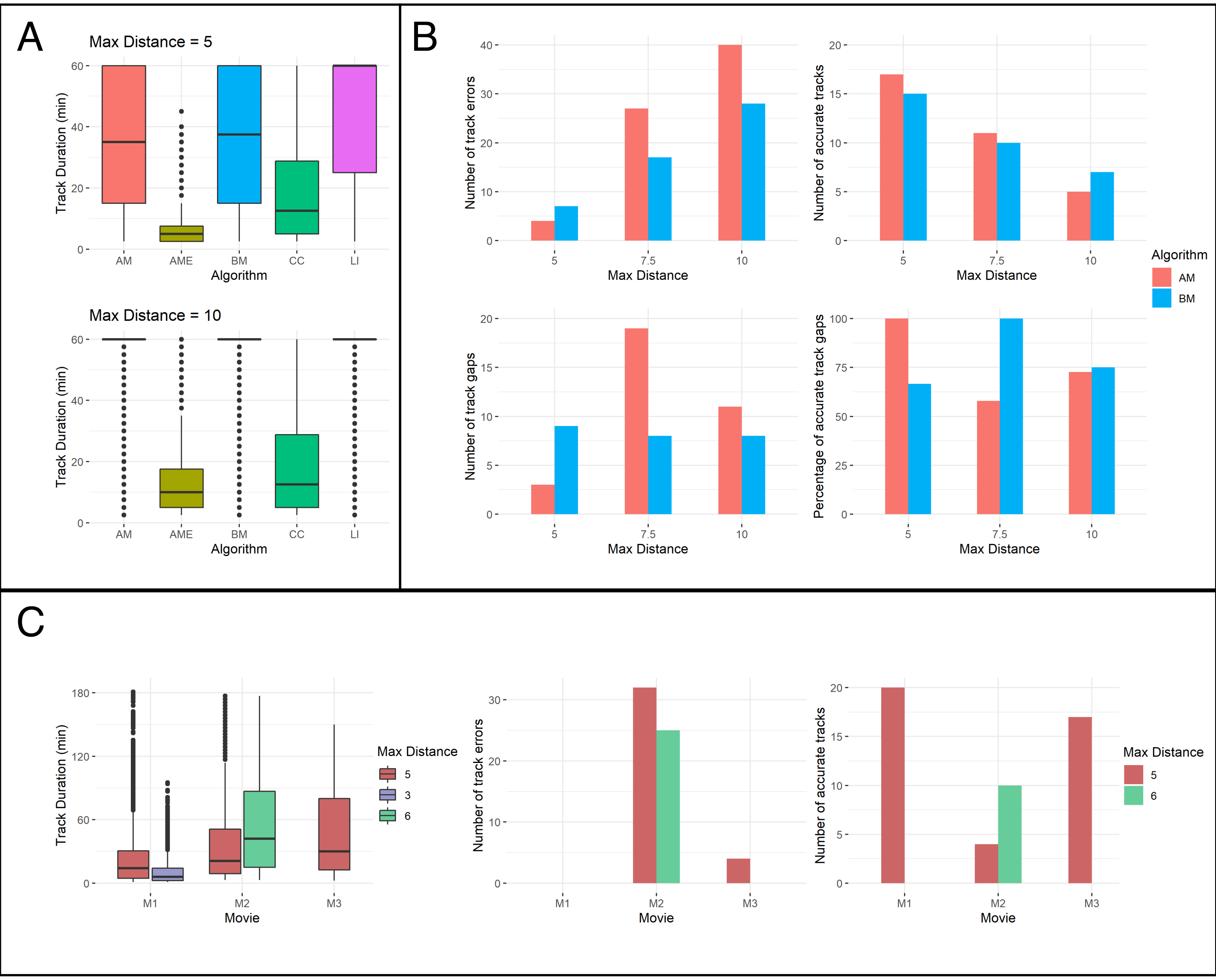

A
Max Distance = 5
Track Duration (min)
Algorithm
AM
AME
BM
CC
LI
Max Distance = 10
B
Number of track errors
Number of accurate tracks
Number of track gaps
Percentage of accurate track gaps
Max Distance
Algorithm
AM
BM
C
Movie
M1
M2
M3
Max Distance
5
3
6


Figure S4

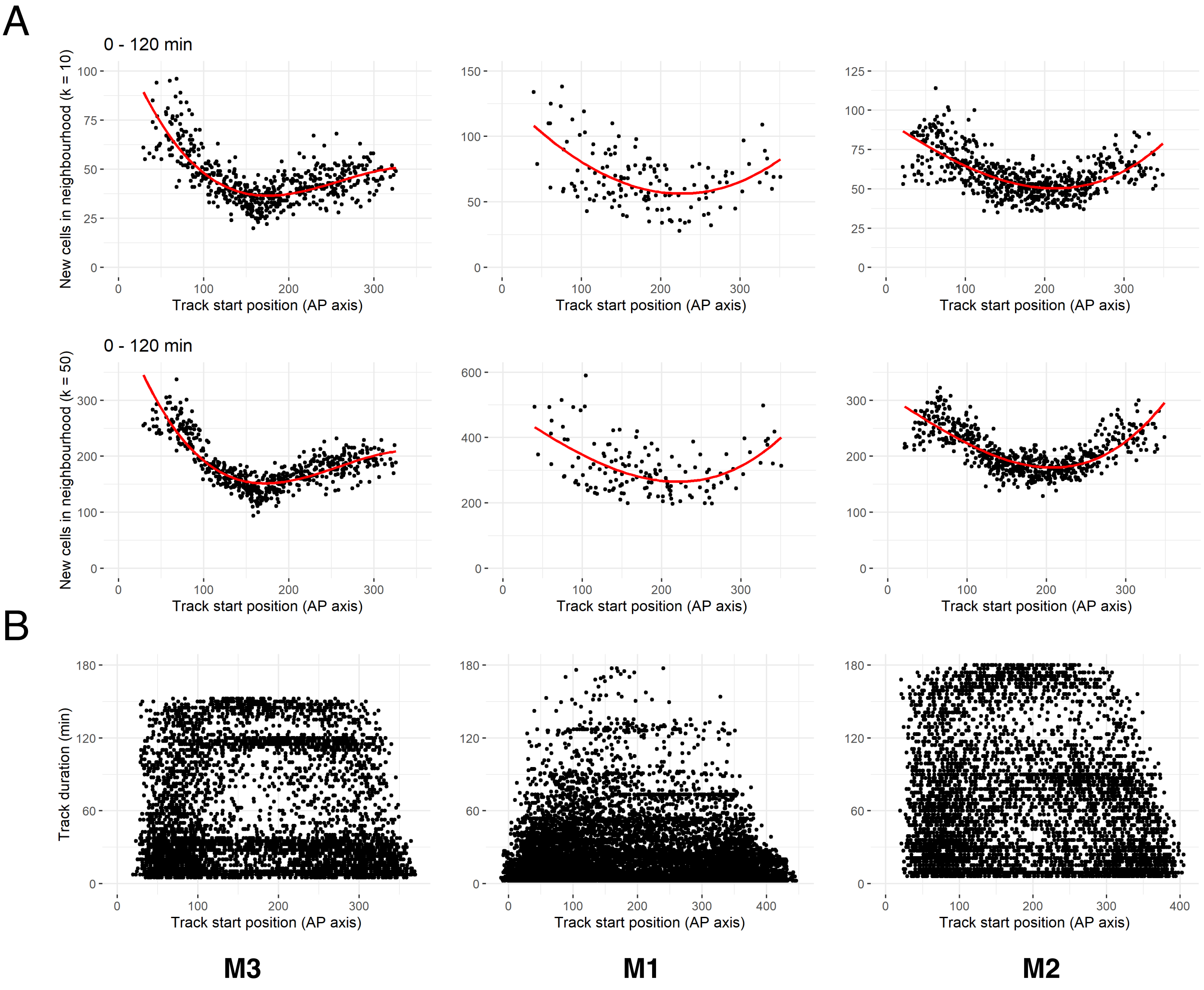

A
0 - 120 min
New cells in neighbourhood (k = 10)
Track start position (AP axis)
0 - 120 min
New cells in neighbourhood (k = 50)
Track start position (AP axis)
B
Track duration (min)
Track start position (AP axis)
M3
M1
M2


Figure S5

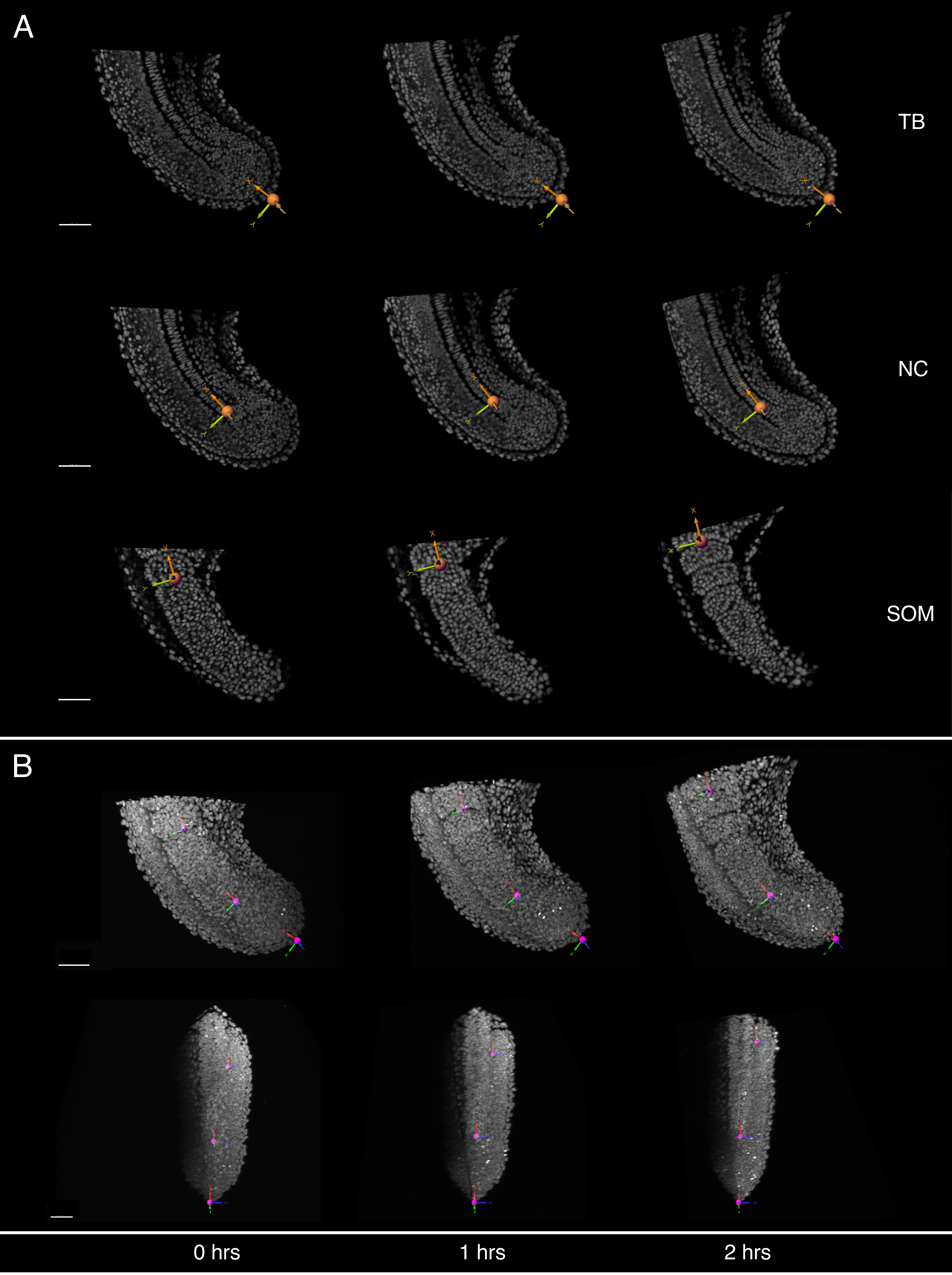


Figure S6

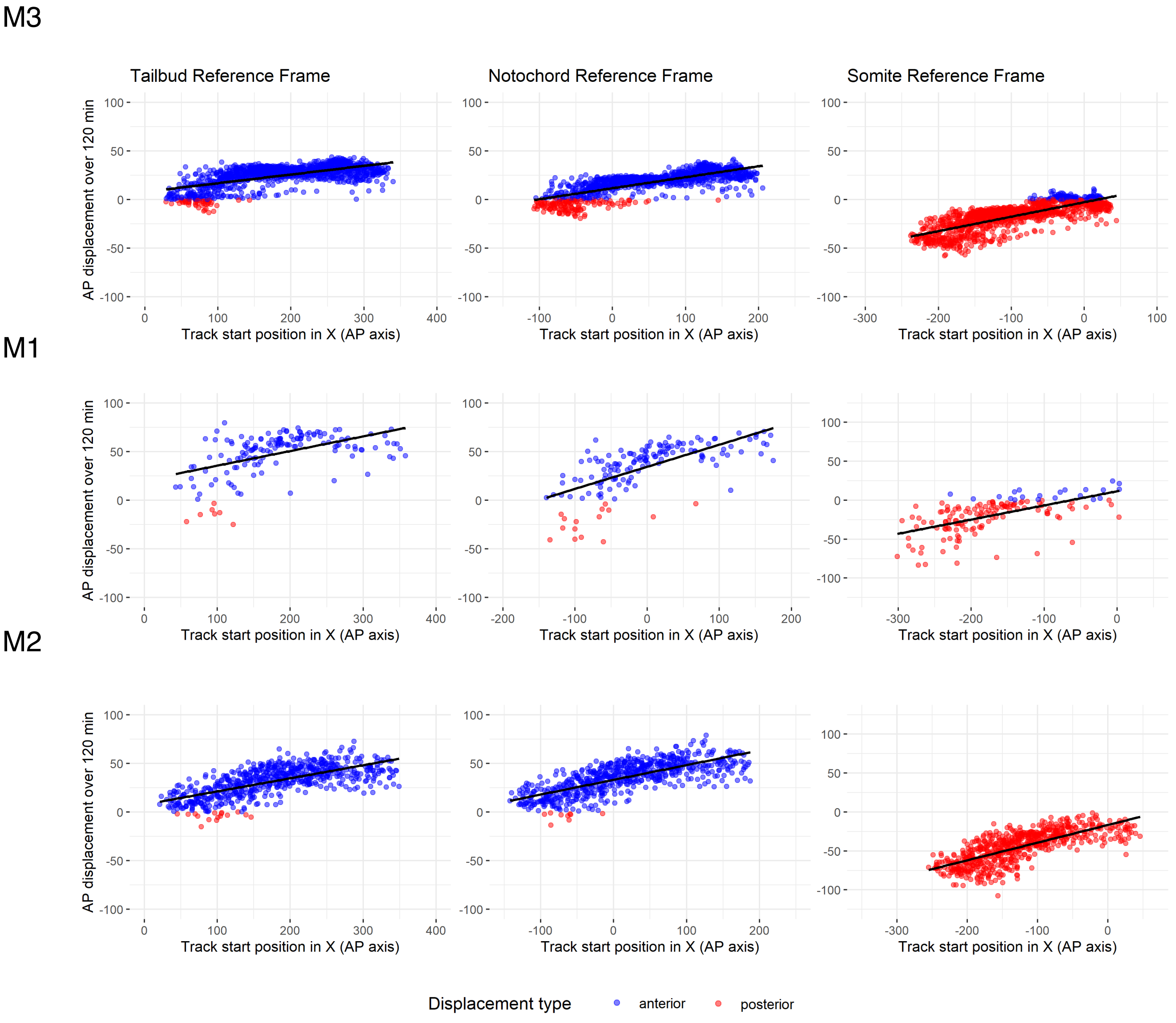
M3
M1
M2
Tailbud Reference Frame
Notochord Reference Frame
Somite Reference Frame
AP displacement over 120 min
Track start position in X (AP axis)
Displacement type
anterior
posterior

Figure S7
